# Generation and characterization of human iPSC-derived *NPC1^I1061T/I10161T^* i^3^Neurons as a model for NPC1 disease

**DOI:** 10.64898/2026.02.11.705111

**Authors:** Shikha Salhotra, Niamh X. Cawley, Christian White, Insung Kang, Anika Prabhu, Cristin D. Davidson, Christopher A. Wassif, Forbes D. Porter

**Author notes:** Corresponding Author: Forbes D. Porter, MD, PhD, 10CRC, Rm. 1-3330, 10 Center Dr, Bethesda, MD 20892. Equivalent contributions (Shikha Salhotra, Niamh X. Cawley). Department of Neuroscience and Pharmacology, Carver College of Medicine, University of Iowa, Iowa City, Iowa, USA.

## Abstract

Niemann-Pick disease, type C is an autosomal recessive, fatal, neurodegenerative disorder caused by pathological variants in *NPC1* or *NPC2*. Dysfunction of either NPC1 or NPC2 results in impaired intracellular cholesterol transport and subsequent storage of unesterified cholesterol in endolysosomal compartments. Earlier cell-based studies utilized patient fibroblasts to study this disease; however, neuronal cells allow for investigation of the neurodegenerative aspect of NPC1. Expression of neurogenin in induced pluripotent stem cells leads to the generation of i^3^Neurons (integrated, isogenic, and inducible), allowing for rapid, synchronized growth of homogenous neurons. In this study, we report the development and characterization of a human iPSC-derived *NPC1^I1061T/I1061T^*i^3^Neuronal model system. *NPC1^I1061T^*is a missense variant resulting in a misfolded protein targeted for proteasomal degradation in the ER. *NPC1^I1061T/I1061T^* i^3^Neurons phenocopied the cellular pathological features of NPC1 disease including endolysosomal cholesterol accumulation, lysosomal morphological changes, and response to the proteostasis modulator, mo56HC. The NPC1 phenotype was alleviated by 2-hydroxypropyl-β-cyclodextrin treatment, a drug demonstrating efficacy both *in vitro* and *in vivo*. This *NPC1^I1061T/I1061T^* i^3^Neuronal cell line can facilitate future high-throughput drug and genomic screens, particularly those aimed at identifying proteostasis regulators that improve the expression/stability of the mutant NPC1 protein.

## 1. Introduction

Cholesterol is a prominent lipid species in the cell that serves critical functions; hence its cellular levels are tightly regulated [1]. Dysregulation of cholesterol homeostasis in the cell can lead to various progressive diseases affecting physical and cognitive abilities. Niemann-Pick disease, type C (NPC), caused by pathogenic variants in *NPC1* or *NPC2*, is one such ultrarare, autosomal recessive, lipid storage disorder. The rate of incidence is approximately 1 in 120,000 live births, with defects in *NPC1* accounting for ∼95% of affected individuals [2]. NPC individuals frequently exhibit hepatosplenomegaly in childhood and manifest insidious, progressive neurodegeneration [2]. NPC1 and NPC2 proteins are cholesterol transporters and play a crucial role in cholesterol egress from lysosomes. Thus, NPC disease is characterized by unesterified cholesterol storage in endolysosomal compartments. The NPC phenotype is heterogeneous with respect to both the age of neurological onset and clinical manifestations. This heterogeneity is likely due to a combination of factors including both residual function of the mutant protein and genetic modifiers [3]. Approximately 200 pathological variants have been identified in *NPC1* [4]. The missense *NPC1^I1061T^*(c. 3182T>C) allele is the most frequent pathological variant found in NPC1 individuals of European heritage. This *NPC1* allele is also common in Hispanic individuals from the Upper Rio Grande Valley in the United States, presumably due to a founder effect [5].

The NPC1 protein has 13 transmembrane domains and a putative sterol sensing domain that is evolutionary conserved and functionally important [6]. The *NPC1^I1061T^* is a transition variant, c.3182T>C, in exon 21 of *NPC1* and was found to influence transmembrane domain 10 of the resultant protein [5]. The NPC1 p.I1061T protein does not efficiently proceed from the endoplasmic reticulum (ER) to the Golgi because it is recognized as a misfolded protein by the ER quality control system and directed for proteasomal degradation [7]. In NPC1-deficient cells, overexpression of *NPC1^I1061T^* results in partial localization of the mutant protein in late endosomes and limited correction of the NPC1 mutant phenotype [7]. Presumably, this is due to some of the nascent NPC1 p.I1061T protein being correctly folded, thereby evading the ER control machinery. Characterization of the murine NPC1 p.I1061T protein in the *Npc1^I1061T/I1061T^*mouse model demonstrated a reduced half-life of the mutant protein [7, 8]. Importantly, when human and murine NPC1 p.I1061T proteins were compared, differences in trafficking through the medial Golgi for each species were observed. Murine protein can traffic from Golgi to late endosomes; however, human NPC1 p.I1061T protein is unable to traffic and remains in the ER. This difference in trafficking pattern has been attributed to variations in protein sequences between the species rather than changes in the ER-folding environment [9]. This finding underscores the need to study the human protein in preclinical models.

Various biological studies and drug screens for treatment of NPC disease have employed fibroblasts from NPC1 individuals [10]. However, given the prominent neurological aspects of NPC1, research needs to be conducted to understand the neuronal pathology of NPC1 disease and to identify potential therapeutics in neurons. We generated neurons from human inducible pluripotent stem cells (iPSCs) containing a system with a doxycycline-inducible neurogenin 2 (*Ngn2*) transgene integrated into the AAVS1 safe-harbor locus. These were originally termed i^3^Neurons to indicate them as isogenic, integrated and inducible [11]. This system is similar to isogenic *NPC1^I1061T^* and other iNeurons generated using the PiggyBac transposase system [9, 12]. The technology allows for the synchronized generation of large numbers of uniform glutamatergic neurons in 10 days. These features make the i^3^Neurons amenable for use in high-throughput drug and genetic screens as well as in molecular and biological studies. This method provides improvements over the standard processes to generate neurons from iPSCs which are generally more complicated and result in a less homogeneous neuronal culture. In addition, i^3^Neurons can be used as control cells in experimental procedures allowing for mutation-specific phenotype characterization.

This study describes the generation and characterization of an iPSC-derived *NPC1^I1061T/I1061T^* i^3^Neuronal line. Our group previously generated and characterized *NPC1^-/-^*i^3^Neurons [13] which are now complemented by this *NPC1^I1061T/I1061T^* i^3^Neuronal model. Research based on the *NPC1^I1061T^* variant is relevant due to its frequency in NPC1 individuals and the possibility of rectifying the error in protein folding via the identification of proteostasis modulators [14]. We show that *NPC1^I1061T/I1061T^*i^3^Neurons have the expected cellular phenotype of NPC1. Specifically, we observed the accumulation of unesterified cholesterol in the endolysosomal system, development of intracellular inclusion bodies, and lysosomal dysfunction. In addition, treatment of these cells with mo56HC, a known proteostasis stabilizer of NPC1, increased levels of the mutant protein. Moreover, 2 hydroxypropyl-beta-cyclodextrin (HPβCD) was able to correct the observed cellular defects in *NPC1^I1061T/I1061T^*i^3^Neurons, thereby providing a paradigm where this hypomorphic model may be useful in high-throughput drug/chemical and genomic screens.

## 2. Materials and Methods

### 2.1 Cell culture and neuronal differentiation

Human iPSCs were grown in Essential 8 Medium (E8) (Gibco/ThermoFisher Scientific, Waltham, MA, #A15169-01) on dishes coated with Matrigel (Corning, Bedford, MA, #354277) according to the manufacturer’s instructions. When the cells reached 70-90% confluency, they were passaged with Accutase (STEMCELL Technologies, Inc, Cambridge, MA, #07920) and cultured in E8 medium supplemented with 10 nM ROCK inhibitor, Y-27632 (Tocris Bioscience/BioTechne, Minneapolis, MN, #1254). The iPSCs were then allowed to grow for 24-48 hours and once colonies were established, fresh E8 media was added. iPSCs were differentiated into i^3^Neurons using the protocol as previously reported [15]. In short, at day 0, iPSCs were cultured in a pre-differentiation medium [Knockout DMEM/F12 medium (Gibco, Waltham, MA, 10829-018) supplemented with 1× GlutaMAX (Gibco, #35050-061), 2 μg/mL doxycycline, 1× MEM non-essential amino acids (Gibco, #11140-050), 1× N2A supplement (STEMCELL Technologies, #07152) and 10 nM ROCK inhibitor]. On days 1 and 2, cultured cells were replenished with the pre-differentiation medium and on day 3, Accutase was used for passaging the cells onto dishes coated with poly-L-ornithine (Sigma-Aldrich, St. Louis, MO, #P3655) in a specific neuronal medium [BrainPhys medium (STEMCELL Technologies, #05790) supplemented with 10 ng/mL NT-3 (PeproTech/ThermoFisher Scientific, #450-03-250 UG), 1 μg/mL mouse laminin (Gibco, #23017-015), 1× NeuroCult SM1 (STEMCELL Technologies #05711) and 10 ng/mL BDNF (PeproTech/ThermoFisher Scientific, #450-02-250 UG)]. To maintain neuronal health, half media changes were done i.e., removal of half media and addition of new media in equal volume, every 2 days.

### 2.2 Generation of *NPC1^I1061T/I1061T^* iPSC line

Isogenic control human iPSCs expressing *Ngn2* in the AAVS1 safe-harbor was originally generated and characterized by Wang et al., [11] and used previously by us for the generation of inducible *NPC1^null^* iPSCs [13]. These control iPSCs were sent to Applied Stem Cell (Milpitas, CA) where an *NPC1^I1061T/I1061T^* iPSC derived neuron-inducible cell line was generated under contract using CRISPR-Cas9 editing technology and homologous recombination with donor templates containing the *NPC1^I1061T^* variant. Two clonal lines were obtained, and the point mutation was confirmed by sequencing.

### 2.3 Genomic DNA extraction, PCR amplification and DNA sequencing

After receipt and expansion of the cells, the sequencing results were verified as follows. Genomic DNA was extracted from both *NPC1^+/+^*and *NPC1^I1061T/I1061T^* iPSCs using a DNA extraction kit (DNeasy Blood and Tissue kit, Qiagen, Germantown, MD, #69506) following the manufacturer’s instructions. The region of the *NPC1* gene around the p.I1061T point mutation was amplified by PCR and sequenced (NICHD Molecular Genomics Core). The sequencing results were analyzed by 4Peaks software (Nucleobytes, https://nucleobytes.com/). The primers used for PCR amplification and sequencing were: Forward: 5’-TGGACTCTCTTGACACCCAG-3’ and Reverse: 5’-ACCCAGTGTAGGCCCTTTG-3’

### 2.4 Western blot of iPSCs

*NPC1^+/+^* and *NPC1^I1061T/I1061T^* iPSCs were plated on Matrigel-coated 6 cm dishes at a density of 250,000 cells/dish. On day 2, cells were lysed in 1x Tissue Protein Extraction reagent (T-PER; ThermoFisher Scientific, #78510) with 1X protease and phosphatase inhibitor cocktail (Sigma, #PPC1010). After 10 minutes, the cell homogenate was collected and centrifuged at 13,000 rpm for 10 min at 4°C. Protein was quantified by the Bradford method using Bio-Rad Protein Assay dye reagent concentrate (Bio-Rad, Hercules, CA, #5000006). For all samples, a total of 20 μg of protein along with pre-stained protein standards (SeeBlue Plus 2 Prestained Standard, Invitrogen/ThermoFisher Scientific, #LC5925) were resolved on a 4–12% Bis/Tris NuPage protein gels (ThermoFisher Scientific, #NP0321BOX) and transferred to a nitrocellulose membrane (iBlot2NC regular stacks, ThermoFisher Scientific, #IB23001) by semi-dry transfer (iBlot2, ThermoFisher Scientific). Membranes were then blocked in 5% non-fat milk diluted in 1X PBST [(1X PBS with 0.01% Tween 20) (1X PBS – Gibco, #10010023, Tween 20 – Sigma-Aldrich, #P7949)] and probed for β-Actin at 1:10,000 (Invitrogen, #MA1-140) and NPC1 at 1:3,000 (Abcam, Waltham, MA, #ab134113).

### 2.5 Endoglycosidase H (Endo H) and PNGase F treatment of iPSCs and i^3^Neurons

*NPC1^+/+^* and *NPC1^I1061T/I1061T^* iPSCs were plated on Matrigel-coated 6 cm dishes at a density of 250,000 cells/dish. On day 2, the cells were harvested with T-PER as described above. To obtain neurons, pre-differentiated *NPC1^+/+^* and *NPC1^I1061T/I1061T^*iPSCs were seeded on day 3 post-doxycycline on poly-L-ornithine coated 10 cm dishes at a density of 3,000,000 cells/dish into the neuronal media. At day 10, cell lysates were made as described above. Deglycosylation reactions were performed for both *NPC1^+/+^*and *NPC1^I1061T/I1061T^* iPSCs and i^3^Neurons with and without the respective enzymes. For both Endo H (New England Biolabs, Ipswich, MA, #P0702S) and PNGase F (New England Biolabs, #P0704S) treatment, 100 μg of total protein for each cell type was combined with 5 μl of 10X glycoprotein denaturing buffer (New England Biolabs, #B1704S) in water to a total volume of 50 μl and then incubated at 100°C for 10 min. For PNGase F treatment, denatured lysate was chilled on ice and centrifuged for 10 seconds. Five μl of 500,000 U/ml PNGase F or an equal volume of water along with 10 μl of 10X Glyco Buffer 2 (New England Biolabs, #B3704S), 10 μl of 10% NP-40 (New England Biolabs, #B2704S) and 1X protease and phosphatase inhibitor cocktail were added to each tube to make a total of 100 μl. These samples were then incubated at 37°C for 1 hour. For Endo H reaction, 5 μl of 500,000 U/ml Endo H or an equal volume of water was added along with 10 μl of 10X Glyco Buffer 3 (New England Biolabs, #B1720S) and 1X protease and phosphatase inhibitor cocktail to make final volume of 100 μl in each tube. Samples were incubated at 37°C for 1 hour. Twenty μl of the samples (control, PNGase-, or EndoH-treated) were then analyzed by western blot as described above.

### 2.6 Immunofluorescence staining for β-III tubulin

*NPC1^+/+^* and *NPC1^I1061T/I1061T^* pre-differentiated i^3^Neurons were plated at day 3 post-doxycycline at a density of 80,000 cells/chamber on poly-L-ornithine coated 2-chamber slides in neuronal media. At day 10, the neurons were washed three times with 1X PBS followed by fixation with 4% paraformaldehyde in 1X PBS [Paraformaldehyde 16% solution, EM grade, Electron Microscopy Sciences (EMS), Hatfield, PA, #15710] for 30 minutes and permeabilized with 0.1% Tween-20 in 1X PBS for 5 minutes. After blocking in 10% normal goat serum (Abcam, #ab7481) in 1X PBS for 1 hour, the cells were then incubated with mouse anti-β-III tubulin primary antibody (R&D Systems, Minneapolis, MN, #MAB1195) at 1:400 dilution overnight at 4°C. After washing three times with 1X PBS, the cells were stained with anti-mouse secondary antibody Alexa Fluor-488 (Invitrogen, #A21202) at 1:1000 dilution for 1 hour. The cells were washed three times with 1X PBS and then cover-slipped using ProLong Gold anti-fade reagent with DAPI (Invitrogen, #P36935) to stain the cell nuclei. The cells were imaged using a Zeiss Axio Observer Z1 inverted fluorescence microscope controlled by ZEN3.1 software (Zeiss, Jena, Germany).

### 2.7 Perfringolysin O (PFO)-iFluor 647 staining of iPSCs and i^3^Neurons

iPSCs from *NPC1^+/+^, NPC1^I1061T/I1061T^* and *NPC1^-/-^* cell lines were plated on Matrigel-coated 2-chamber slides at a density of 35,000 cells/chamber. i^3^Neurons from all three cell lines were seeded at day 3 post-doxycycline on poly-L-ornithine coated 2-chamber slides in neuronal media at a density of 80,000 cells/chamber. Day 2 iPSCs and day 10 i^3^Neurons were fixed, permeabilized and blocked as described for immunofluorescence and then incubated with PFO-iFluor 647 (Codex Biosciences/Dexorgen, Rockville, MD, #CB-170411c) at 1.7 μg/ml for 1 hour. The cells were then washed three times with 1X PBS, and the nuclei were stained using ProLong Gold anti-fade reagent with DAPI, cover-slipped and visualized as described above.

### 2.8 Co-localization of LysoTracker Red and PFO-Alexa Fluor-488

*NPC1^+^*^/+^, *NPC1^I1061T/I1061T^*and *NPC1^-/-^* pre-differentiated i^3^Neurons were plated at day 3 post-doxycycline on poly-L-ornithine coated 12-well plates at a density of 100,000 cells/well in neuronal media. On day 10, LysoTracker Red (LysoTracker™ Red DND-99, Invitrogen, #L7528) was used at final concentration of 100 nM in neuronal media for 2 hours at 37°C. Cells were washed, fixed, and permeabilized as described above. The cells were then stained with PFO-Alexa Fluor-488 at 0.89 μg/ml for 1 hour. The cells were washed three times with 1X PBS and stained using Hoechst (Hoechst 33342, ThermoFisher Scientific, #H3570) at 1:10,000 dilution in 1X PBS for 10 minutes to visualize the cell nuclei. After rinsing with 1X PBS, the cells were imaged using the Zeiss Axio Observer Z1 microscope described above.

### 2.9 Electron Microscopy

Pre-differentiated day 3 i^3^Neurons for both *NPC1^+/+^*and *NPC1^I1061T/I1061T^* were plated on MatTek glass dishes (35 mm dish, No.1.5 Coverslip, 14 mm glass diameter, uncoated, MatTek Life Sciences, Ashland, MA, #P35G-1.5-14-C) at a density of 90,000 cells/dish in BrainPhys medium. On day 10, the media was removed, and neurons were washed gently 1-3 times in 1X PBS and fixed using room temperature (RT) EM fixative [2.5% glutaraldehyde made in 0.1M sodium cacodylate buffer, pH 7.4) (Aqueous Glutaraldehyde EM Grade 25%, EMS, #16220) (Sodium Cacodylate buffer, 0.2 M, pH 7.4, EMS, #11652)]. Cells were fixed at RT for 30 minutes with gentle agitation on a rocker. The fixative was removed, and cells washed 3 times with EM buffer (0.1M sodium cacodylate buffer, pH 7.4) for 5 minutes each. Using the variable wattage Pelco BioWave Pro microwave oven (Ted Pella, Inc., Redding, CA), the subsequent processing steps were performed: 0.1M sodium cacodylate buffer, pH 7.4 was used to wash the cells followed by additional fixation with 1% osmium tetroxide (EMS, #19100) in 0.1M sodium cacodylate buffer for 10 minutes. Cells were then washed in double-distilled water (DDW) followed by enhancement with 3% (aqueous) uranyl acetate (EMS, #22400-3) for 60 minutes at room temperature on a rocker. The cells were washed again with DDW and then dehydrated by sequentially increasing ethanol to 100% followed by the ultimate infiltration by an Embed-812 resin (EMS, #14900) which was sequentially increased to 100%. Polymerization of resin was achieved by incubating samples in an oven set at 60°C for 20 h. A Leica EM UC7 ultramicrotome (Leica Biosystems, Nussloch, Germany) was used to generate 90 nm ultra-thin sections. Thin sections were placed on 200 mesh copper grids and post-stained with UranyLess (Uranyl Acetate substitute, EMS, #22409) and lead citrate. Imaging was performed on a JEOL-1400 Transmission Electron Microscope operating at 80kV and an AMT BioSprint-29 camera (NICHD Microscopy Core Facility).

### 2.10 Treatment of i^3^Neurons with potential therapeutic compounds

Pre-differentiated i^3^Neurons from *NPC1^+/+^*, *NPC1^I1061T/I1061T^* and *NPC1^-/-^* iPSCs were plated on poly-L-ornithine coated 2-chamber slides at day 3 post-doxycycline at a density of 80,000 cells/chamber in neuronal media. At day 10, i^3^Neurons were treated overnight with 100 µM 2-hydroxypropyl-β-cyclodextrin (HPβCD; Kleptose® HPB, Roquette, Geneva, IL). Processing and visualization of the cells using PFO-647 were as described in the section on PFO staining.

To stabilize the *NPC1^I1061T/I1061T^* mutant protein, mo56HC was used (graciously provided by Dr. Brian Blagg at the University of Notre Dame (Notre Dame, IN)). *NPC1^+/+^* and *NPC1^I1061T/I1061T^*pre-differentiated i^3^Neurons were seeded on poly-L-ornithine coated 10 cm dishes at day 3 post-doxycycline at 3,000,000 cells/dish in neuronal media. i^3^Neurons at day 10 were treated with 10 μM mo56HC for 48 hours at 37°C. The cells were then harvested and lysed followed by determination of protein concentration and western blotting for NPC1, as described above.

### 2.11 Quantitation methods and statistical analysis

ImageJ 1.53t was used to analyze the microscope images for puncta count and puncta area, which were normalized to nuclear count. For colocalization analysis, Pearson’s colocalization coefficient was calculated using the ImageJ Plugin, JaCoP. The results are expressed as the mean ± SD of three independent experiments, wherein 100 cells were counted per cell line for each experiment. Comparison between the two groups was performed by unpaired t-test. Statistical analyses were performed using GraphPad Prism version 9.5.1 (GraphPad Software, San Diego, CA, USA).

## 3. Results

### 3.1 Generation and confirmation of *NPC1*^I1061T^ mutation

CRISPR-Cas9 editing was used to introduce the genomic DNA change- Chr18:23536736 (on Assembly GRCh38) [c.3182T>C] variant into the control iPSCs [16]. To confirm the genotype, genomic DNA was sequenced by Sanger sequencing and the T>C transition in the DNA was verified (**Fig. 1A, B** and **Fig. S1**). Analysis of the sequencing chromatograms also demonstrated the *NPC1* c.3182T>C variant to be homozygous as shown by a single peak. Additionally, we observed the presence of a heterozygous *NPC1* variant c.3174C>A which results in a silent change at NPC1 p.A1058A.

**Figure 1.**
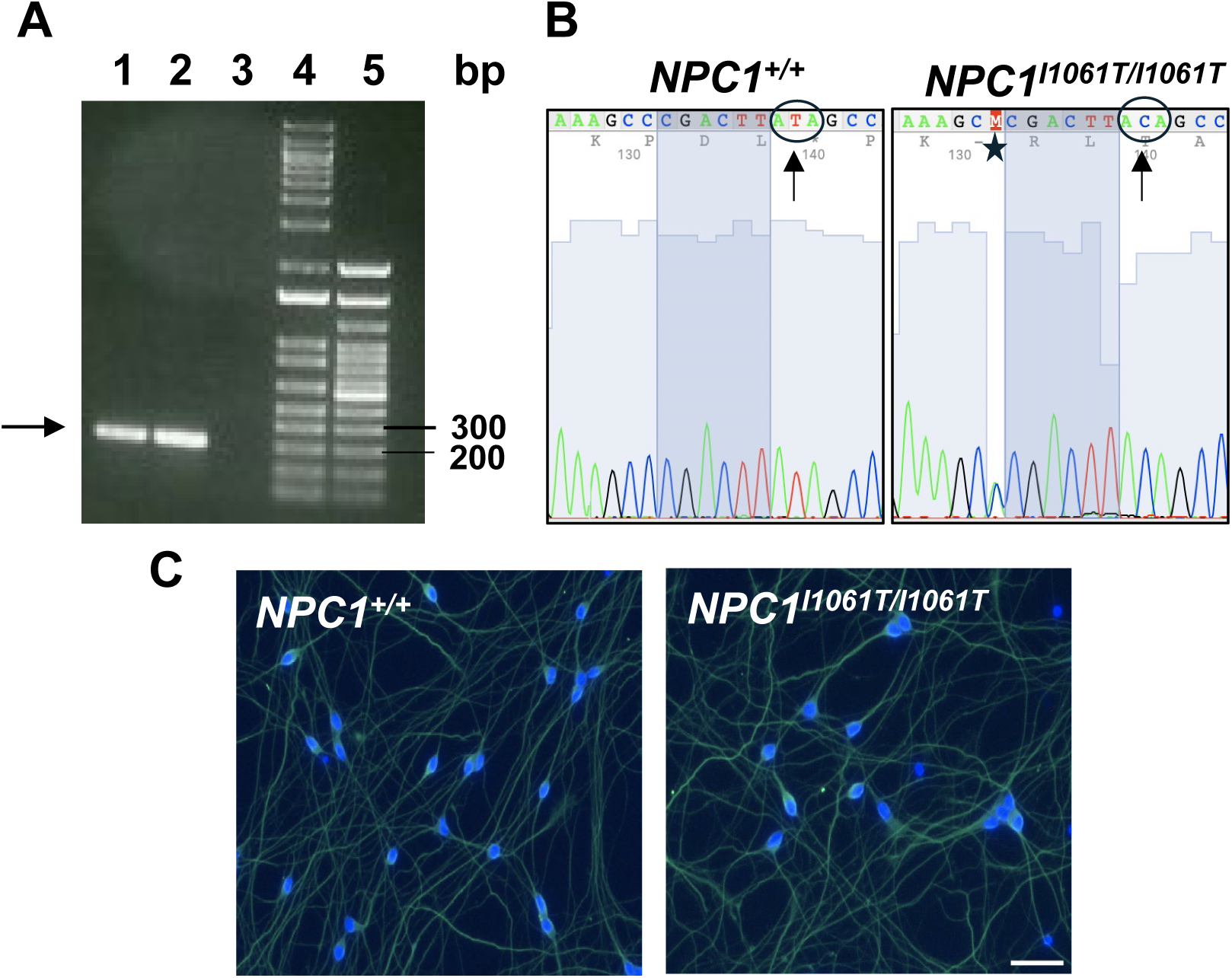
Confirmation of *NPC1^I1061T^* mutation and differentiation. (**A**) Lanes 1 and 2 represent the expected PCR product (arrow) from *NPC1^+/+^* and *NPC1^I1061T/I1061T^* iPSCs, respectively. Lane 3: Non-template control; Lanes 4 and 5: 1 kb and 100 bp DNA ladders, respectively. (**B**) Forward sequencing chromatograms of the PCR products obtained in (A). Note the T to C transition (arrows) at position 3182 resulting in isoleucine substitution by threonine. Sequencing in the reverse direction confirmed this result (**Fig. S1**). Of note was the identification of an additional silent mutation c.3174C>A (star), (Ala to Ala). (**C**) β-III tubulin (neuronal stain, green) and DAPI (nuclear stain, blue) immuno-fluorescence of fully differentiated *NPC1^+/+^*and *NPC1^I1061T/I1061T^* i^3^Neurons (scale bar, 50 μm).

To confirm that the generated iPSCs maintained their stem properties during the CRISPR-Cas9 editing, the cells were differentiated into i^3^Neurons and stained for β−III tubulin, a widely recognized marker of neuronal differentiation [17]. Both *NPC1^I1061T/I1061T^* and *NPC1^+/+^* i^3^Neurons were uniformly stained for β-III tubulin demonstrating that iPSCs completely differentiated into i^3^Neurons in both genotypes (**Fig. 1C**).

### 3.2 NPC1 p.I1061T is sensitive to Endoglycosidase H digestion

Previous studies have shown that NPC1 p.I1061T protein undergoes proteasomal degradation due to protein misfolding in the ER [7]. Endoglycosidase H (Endo H) is an enzyme that catalyzes the removal of N-linked glycans from immature glycoproteins. Proteins that exhibit impaired transit from the ER to the Golgi are sensitive to Endo H digestion. However, if the protein has translocated from the ER to the Golgi, it becomes Endo H-resistant having undergone complex N-linked glycosylation in the Golgi [7]. As a positive control, we used PNGase F, an enzyme which removes all N-linked glycans from glycoproteins, regardless of complexity. Our results demonstrate that the NPC1^I1061T^ protein from *NPC1^I1061T/I1061T^* iPSCs (**Fig. 2A**) and i^3^Neurons (**Fig. 2B**) is more sensitive to Endo H digestion compared to NPC1 from *NPC1^+/+^* iPSCs and i^3^Neurons, respectively. As expected, NPC1 protein from both *NPC1^+/+^* and *NPC1^I1061T/I1061T^* iPSCs and i^3^Neurons were sensitive to PNGase F digestion (**Fig. 2A, B**).

**Figure 2.**
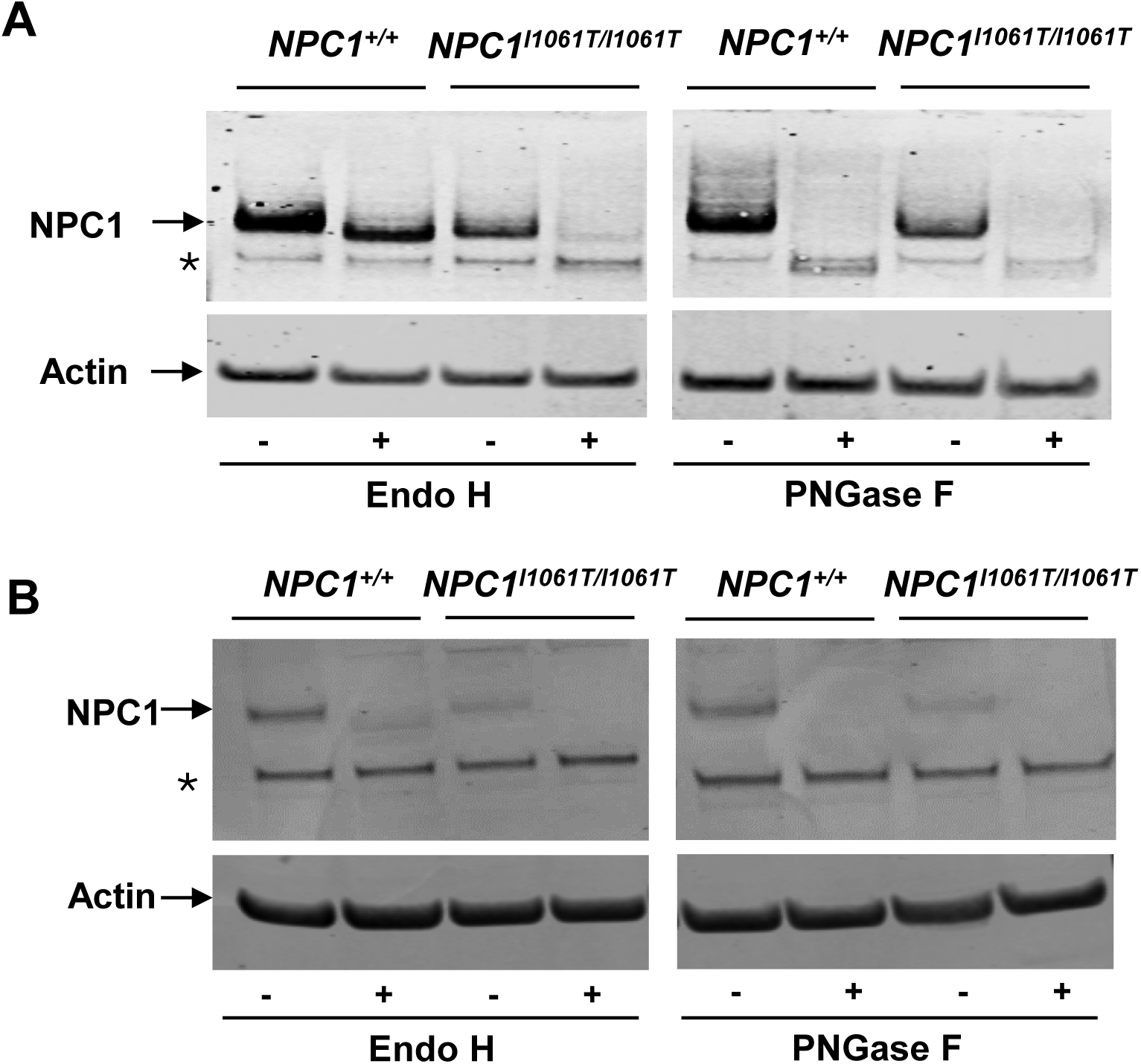
NPC1^I1061T^ mutant protein is sensitive to Endo H in both iPSCs and i^3^Neurons. Western blot analysis of NPC1 in iPSC- (**A**) or i^3^Neuron- (**B**) extracts with and without Endo H (left panels) and PNGase F (right panels) digestion. NPC1^I1061T/I1061T^ is sensitive to both Endo H and PNGase F. NPC1^+/+^ is resistant to Endo H but sensitive to PNGase F. Data is representative of 3 independent experiments. * represents non-specific band.

### 3.3 Endolysosomal accumulation of unesterified cholesterol

Accumulation of unesterified cholesterol in acidic compartments of the cell is a characteristic feature of NPC1-deficient cells [2]. To detect unesterified cholesterol, we used Perfringolysin-O conjugated to iFluor-647 (PFO-647) on iPSCs and i^3^Neurons from three different cell lines: *NPC1^+/+^*, *NPC1^I1061T/I1061T^* and *NPC1*^-/-^ to investigate cholesterol accumulation [18]. *NPC1^-/-^*cells were included as a positive control, as described previously [13]. Our results here showed low-level staining of PFO in *NPC1^+/+^* cells in both iPSCs and i^3^Neurons (**Fig. 3A, B**). However, *NPC1^I1061T/I1061T^* iPSCs and i^3^Neurons showed higher PFO puncta count and puncta size than *NPC1^+/+^* cells (**Fig. 3A, B**). The number of PFO puncta per cell was 2.4-fold higher (p<0.01) in *NPC1^I1061T/I1061T^* iPSCs and 2.3-fold higher (p<0.05) in *NPC1^I1061T/I1061T^* i^3^Neurons as compared to *NPC1^+/+^* iPSCs and i^3^Neurons, respectively. Similarly, the size of puncta (μm^2^) per cell increased by 5.2-fold (p<0.01) in *NPC1^I1061T/I1061T^* iPSCs and 5.6-fold (p<0.001) in *NPC1^I1061T/I1061T^* i^3^Neurons when compared to *NPC1^+/+^* iPSCs and i^3^Neurons, respectively.

**Figure 3.**
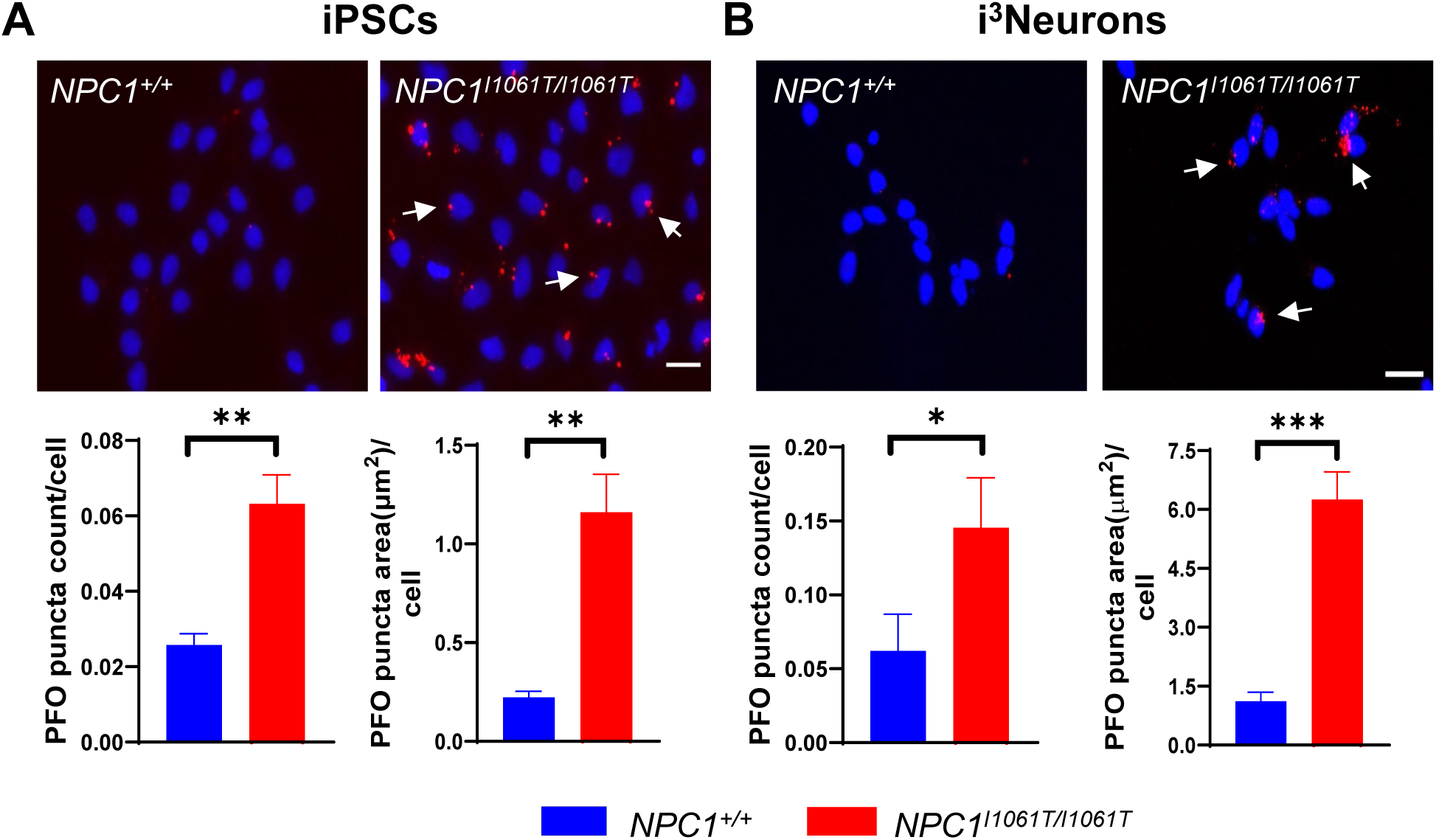
Characterization of cholesterol accumulation in lysosomes using PFO staining. Perfringolysin O-iFluor 647 (red) was used to observe unesterified cholesterol accumulation, (white arrows) in iPSCs (**A**) and i^3^Neurons (**B**). *NPC1^I1061T/I1061T^* iPSCs and i^3^Neurons have higher PFO puncta count as well as puncta area per cell compared to *NPC1^+/+^*control cells (bottom panels, n=3). Nuclei were counterstained with DAPI (blue). Scale bar, 20 μm.*p<0.05, **p<0.01, ***p<0.001, unpaired t-test.

As expected, *NPC1^-/-^* iPSCs and i^3^Neurons showed more prominent cholesterol storage via PFO labeling due to the more severe nature of the mutation (i.e. a null mutation vs missense I1061T). Both the size of PFO puncta as well as puncta count were increased significantly in *NPC1^-/-^* cells as compared to *NPC1^+/+^* cells for both iPSCs and i^3^Neurons, respectively (**Fig. S2**).

### 3.4 Co-localization of LysoTracker Red and PFO-488

In NPC1 disease, cholesterol accumulation is known to occur in the late endosome/lysosomal compartments of the cell [2]. To verify whether cholesterol is accumulating in these organelles in *NPC1^I1061T/I1061T^* i^3^Neurons, LysoTracker™ Red, a viable fluorescent probe that preferentially stains acidic compartments such as late endosomes/lysosomes, was used in conjunction with PFO-488 staining [19]. *NPC1^+/+^* i^3^Neurons had no detectable co-staining with LysoTracker Red as they have very little PFO staining. Our results demonstrated co-staining of LysoTracker Red with the PFO signal in specific regions of *NPC1^I1061T/I1061T^* i^3^Neurons, indicating cholesterol accumulation in the endosomal/lysosomal system (**Fig. 4A, arrows**). When quantified, *NPC1^I1061T/I1061T^* i^3^Neurons showed higher PFO puncta count and puncta size along with increased lysotracker red puncta count and puncta area as compared to *NPC1^+/+^* i^3^Neurons (**Fig. 4B**). The size of PFO puncta (μm^2^) per cell and the number of PFO puncta per cell increased by 5.3-fold (p<0.01) and 4.6-fold (p<0.01) in *NPC1^I1061T/I1061T^* i^3^Neurons as compared to *NPC1^+/+^* i^3^Neurons, respectively. Similarly, the lysotracker red puncta count per cell was 3.95-fold higher (p<0.01), and the lysotracker red puncta size (μm^2^) per cell was 5.7-fold higher (p<0.05) in *NPC1^I1061T/I1061T^* i^3^Neurons when compared to *NPC1^+/+^* i^3^Neurons, respectively. Pearson’s colocalization coefficient which is a measure of the degree of overlap between two channels, was calculated and found to be 11.4-fold (p<0.0001) higher in *NPC1^I1061T/I1061T^* i^3^Neurons, confirming increased co-localization in these cells as compared to *NPC1^+/+^* i^3^Neurons (**Fig. 4B**).

**Figure 4.**
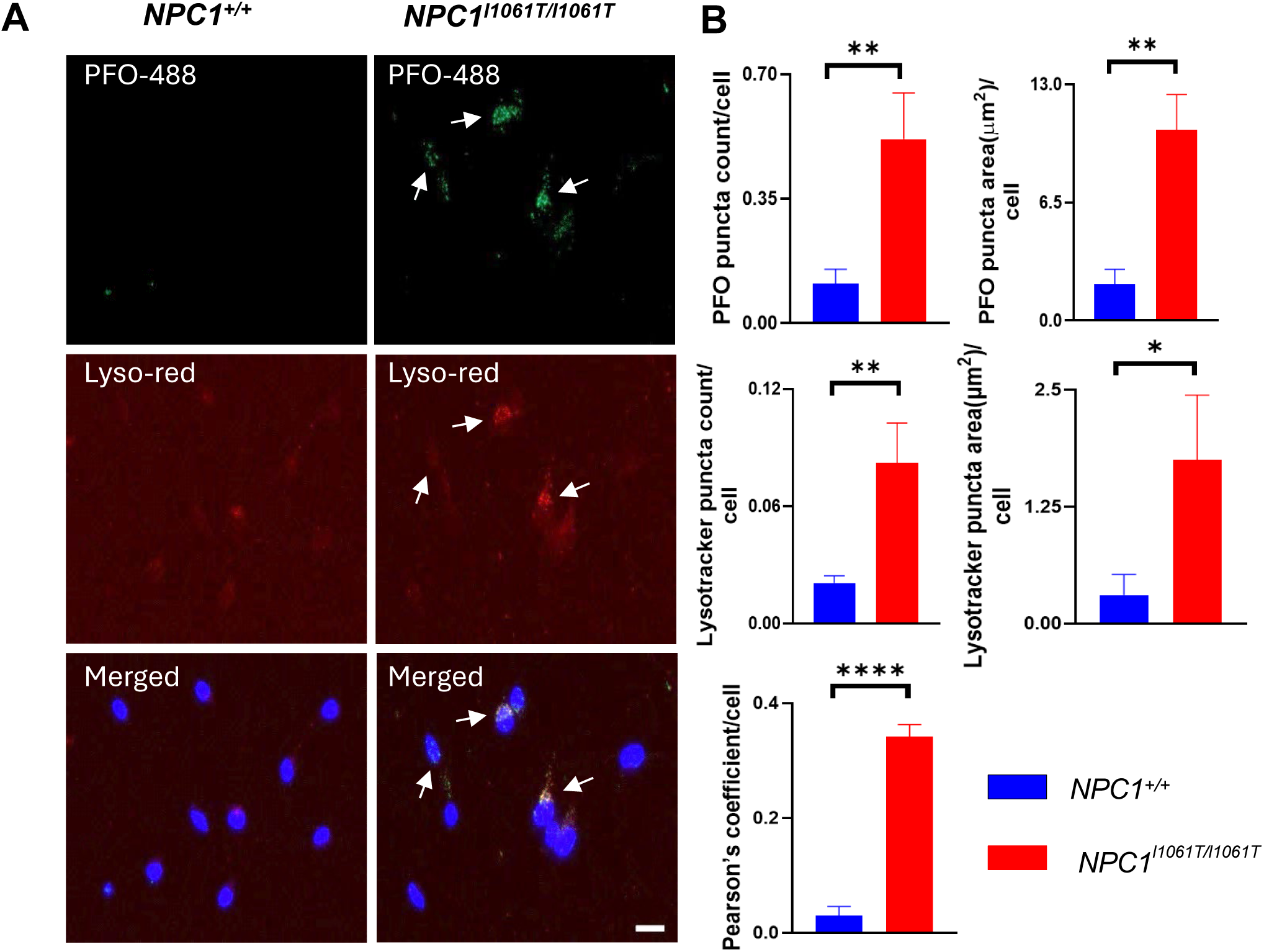
Endolysosomal accumulation of cholesterol in *NPC1^I1061T/I1061T^* i^3^Neurons. **(A)** Co-staining of PFO-488 (green) with LysoTracker Red indicates accumulated cholesterol present in the endo-lysosomal compartment (arrows). Co-staining was higher in *NPC1^I1061T/I1061T^* i^3^Neurons as compared to *NPC1^+/+^*. **(B)** *NPC1^I1061T/I1061T^* i^3^Neurons have higher PFO puncta area (μm^2^) and puncta count per cell as well as increased lysotracker red puncta count and puncta area per cell compared to *NPC1^+/+^*. Pearson’s coefficient was found to be higher in *NPC1^I1061T/I1061T^* i^3^Neurons in comparison to *NPC1^+/+^*. Nuclei were counterstained with DAPI (blue). Scale bar, 20 μm. *p<0.05, **p<0.01, ****p<0.0001 using unpaired t-test.

A similar co-localization was observed in the *NPC1^-/-^*i^3^Neurons. Both PFO puncta count and puncta size per cell as well as lysotracker red puncta count and puncta size per cell were found to be significantly higher in *NPC1^-/-^*i^3^Neurons as compared to *NPC1^+/+^* i^3^Neurons, respectively. Pearson’s colocalization coefficient in *NPC1^-/-^* confirmed the expected cholesterol accumulation in acidic vesicles of this previously characterized model system (**Fig. S3**).

### 3.5 Visualization of storage bodies using electron microscopy

Cholesterol accumulation in lysosomes leads to morphological changes in these organelles. Electron microscopy images from both *NPC1^+/+^* and *NPC1^I1061T/I1061T^* i^3^Neurons showed dark, compact lysosomes. However, lysosomes from *NPC1^I1061T/I1061T^*i^3^Neurons also displayed a distinct pattern of multi-lamellar inclusion bodies (**Fig. 5, inset**), previously reported in NPC1 disease [20]. These results are consistent with endolysosomal storage of unesterified cholesterol in *NPC1^I1061T/I1061T^* i^3^Neurons.

**Figure 5.**
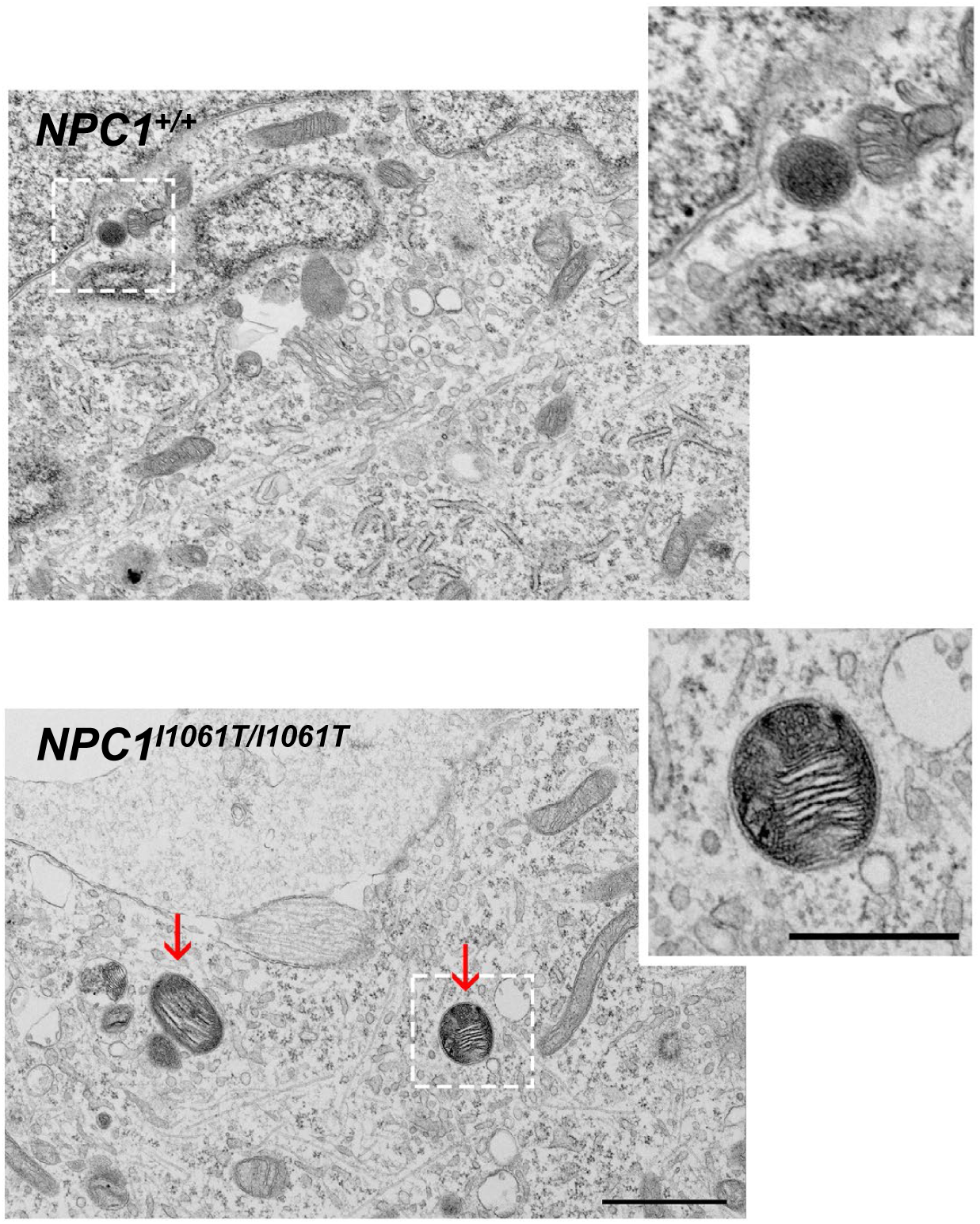
*NPC1^I1061T/I1061T^* i^3^Neurons show morphological changes in lysosomes. i^3^Neurons from *NPC1^I1061T/I1061T^*showed electron-dense, multi-lamellar structures in EM analyses, while *NPC1^+/+^*i^3^Neurons had electron-dense bodies without pathological changes. Insets highlight a normal (upper) and abnormal (lower) lysosome. Scale bar, 1 μm for the image and 0.5 μm for the inset. Arrows indicate multi-lamellar lysosomes in *NPC1^I1061T/I1061T^* i^3^Neurons.

### 3.6 Reduction of PFO staining by 2-hydroxypropyl-β-cyclodextrin (HPβCD)

HPβCD, a compound known to increase cholesterol solubility, reduces the levels of unesterified cholesterol in NPC1 cell lines [21, 22] and has shown efficacy in treating NPC1 disease in animal models and NPC1 individuals [23–26]. In our study, we showed by fluorescence microscopy a decrease of accumulated cholesterol in the lysosomes of *NPC1^I1061T/I1061T^* i^3^Neurons following HPβCD treatment (**Fig. 6A).** When quantified, we found PFO puncta count per cell and puncta area per cell decreased by 2.6-fold (p<0.05) and 3.5-fold (p<0.01), respectively, for *NPC1^I1061T/I1061T^* i^3^Neurons (**Fig. 6B**). A more robust result was observed for the control *NPC1^-/-^*cells as expected (**Fig. S4**). Similar reductions were seen in the *NPC1^+/+^*i^3^Neurons treated with HPβCD where puncta area per cell and puncta count per cell decreased by 4.2 fold (p<0.05) and 2.9 fold (p<0.01), respectively.

**Figure 6.**
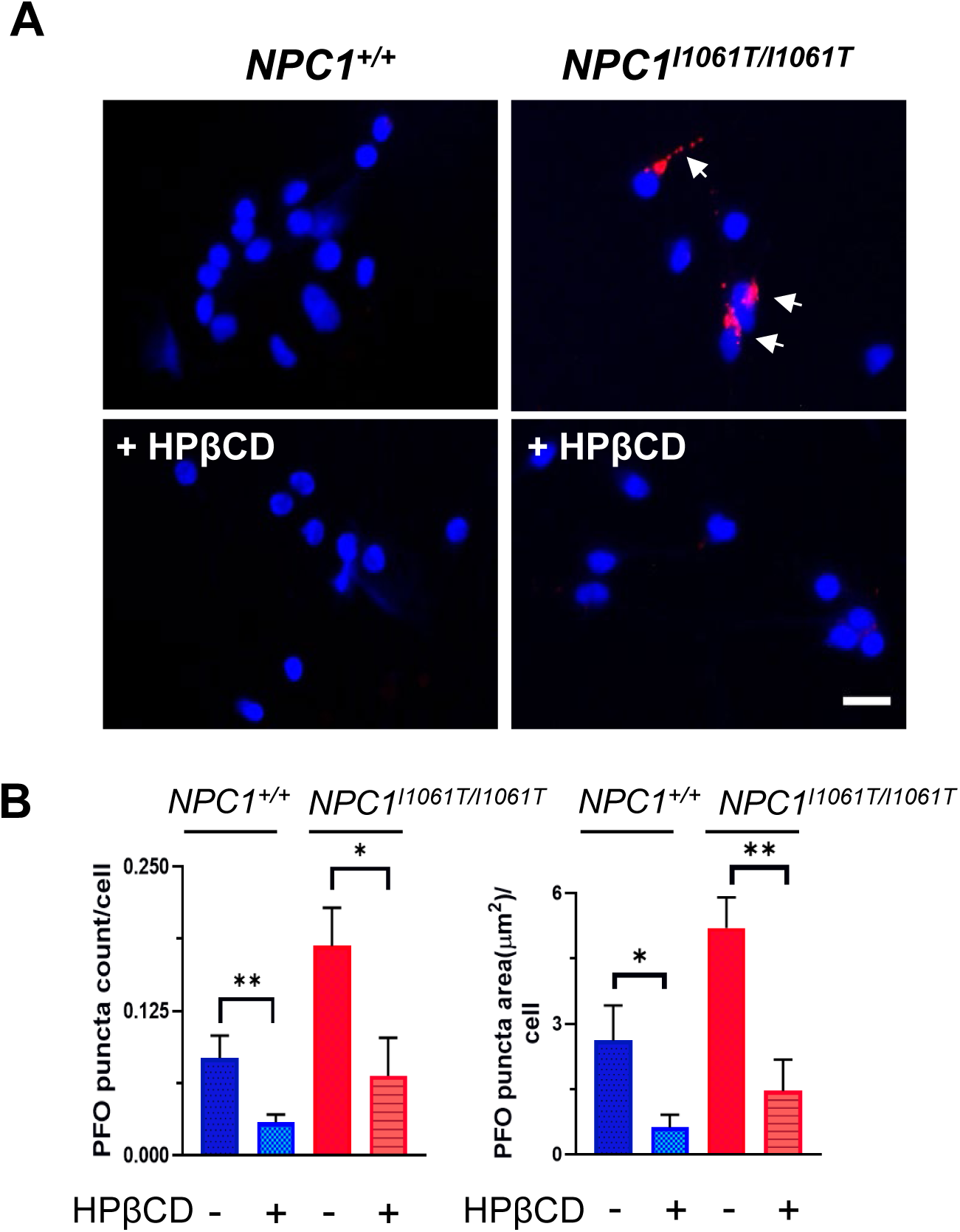
2-Hydroxypropyl-β-cyclodextrin (HPβCD) reduces PFO staining in *NPC1^I1061T/I1061T^* i^3^Neurons. (**A**) PFO-647 staining (red) of *NPC1^+/+^* and *NPC1^I1061T/I1061T^*i^3^Neurons treated with 100 μM of HPβCD. **(B)** PFO puncta count and puncta size (μm^2^) per cell were reduced after HPβCD treatment in both *NPC1^+/+^* and *NPC1^I1061T/I1061T^* i^3^Neurons. Nuclei were counterstained with DAPI (blue). Scale bar, 20 μm. *p<0.05, **p<0.01 using unpaired t-test.

### 3.7 Stabilization of NPC1 p.I1061T in i^3^Neurons by mo56HC

mo56HC is an oxysterol derivative which has been shown to act as a pharmacological chaperone by binding to and stabilizing the mutant NPC1^I1061T^ protein to overcome the trafficking defect [27]. Here, we investigated whether mo56HC could rescue the levels of NPC1^I1061T^ protein in our *NPC1^I1061T/I1061T^*i^3^Neurons. When *NPC1^I1061T/I1061T^* i^3^Neurons were treated with 10 μM mo56HC for 48 hours, NPC1 protein levels increased with a slight shift in band size comparable to the NPC1 protein size from *NPC1^+/+^* i^3^Neurons (**Fig. 7A)**. Quantification of the western blots showed that the NPC1^I1061T^ protein levels increased by 1.4-fold (p<0.05) in *NPC1^I1061T/I1061T^*i^3^Neurons after mo56HC treatment (**Fig. 7B)**.

**Figure 7.**
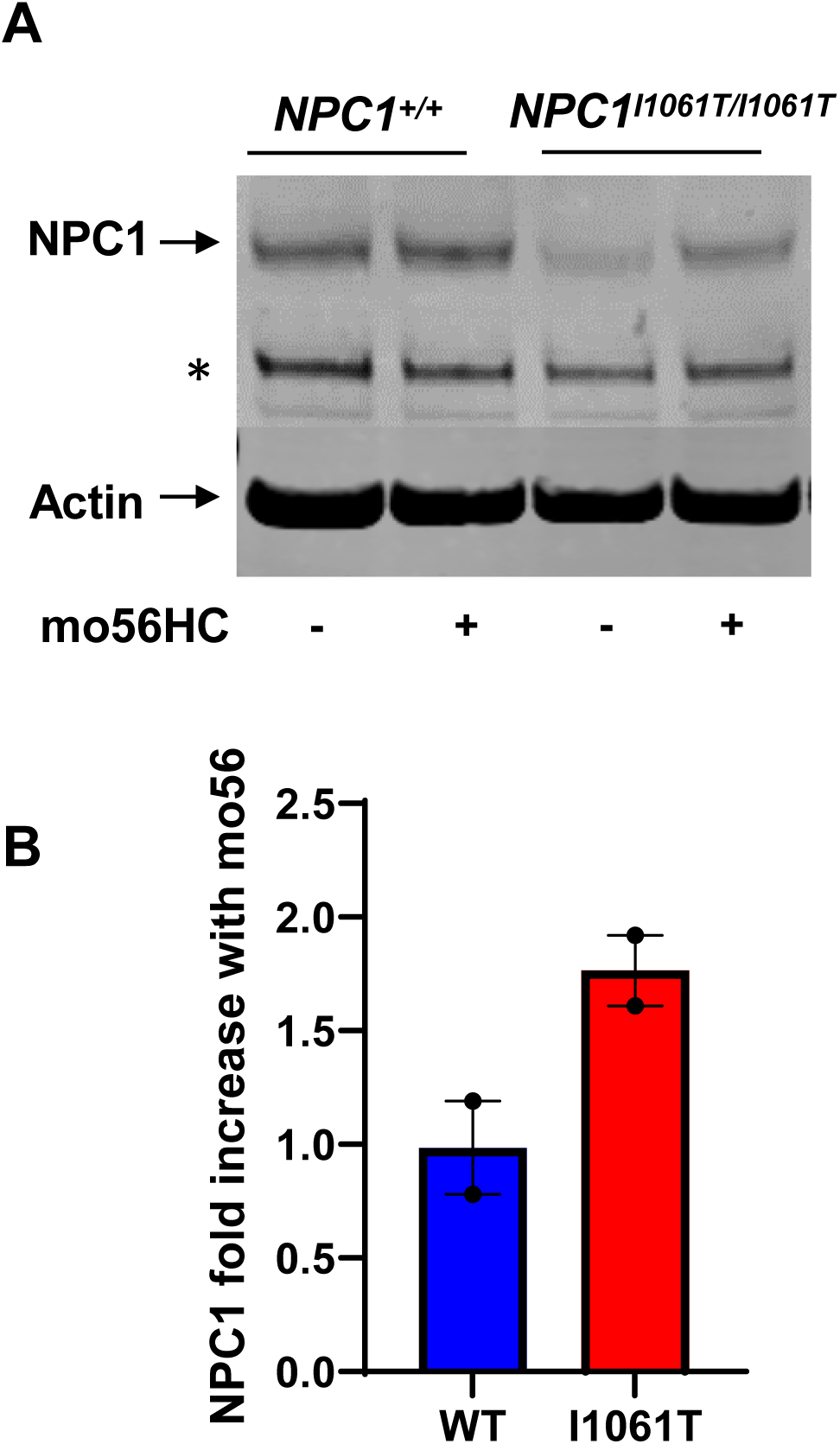
Stabilization of NPC1 p.I1061T protein by mo56HC. **(A)** Western blot analysis of NPC1 from *NPC1^+/+^* and *NPC1^I1061T/I1061T^*i^3^Neurons treated with 10 μM of mo56HC. * non-specific band. **(B)** Average fold change of NPC1 protein normalized to β-actin and control for *NPC1^+/+^* and *NPC1^I1061T/I1061T^* i^3^Neurons, respectively. NPC1^I1061T^ protein increased by 1.8-fold upon treatment with mo56HC. NPC1 protein in *NPC1^+/+^* i^3^Neurons showed no observable change. Data collected from 2 biological replicates.

## 4. Discussion

This study reports the development and characterization of a hypomorphic *NPC1^I1061T/I1061T^* iPSC line with integration of an inducible *NGN2* transgene. Doxycycline-induced neurogenin expression allows for the rapid, large-scale, synchronous and homogeneous differentiation into neurons. These i^3^Neurons are well-suited for high-throughput genetic and small molecule screens. The *NPC1^I1061T/I1061T^* i^3^Neuronal line described here manifests an intermediate NPC1 cellular phenotype and thus complements our previously reported *NPC1^-/-^*i^3^Neuronal line. Given that the NPC1 p.I1061T protein is still functional even though it undergoes intracellular degradation due to misfolding [7], this *NPC1^I1061T/I1061T^*i^3^Neuron line may be particularly useful in screening and testing potential proteostatic regulators. Indeed, a recent similar *Ngn2* inducible iPSC line has been reported and found to fully conform with the expected phenotype of *NPC1^I1061T/I1061T^*neuronal cells [12].

The *NPC1^I1061T/I1061T^* i^3^Neuronal model displays the expected cellular phenotype of NPC1 disease and therefore provides a relevant model for studying missense variants. Moreover, the ability to correct the protein misfolding via modulation of chaperone machinery provides a logical therapeutic approach [14]. *NPC1^I1061T/I1061T^* i^3^Neurons provide a system similar to *NPC1^-/-^*i^3^Neurons in which there is an accumulation of unesterified cholesterol in the endolysosomal compartment of the cell [13], demonstrated by co-staining of PFO and LysoTracker Red. However, as anticipated, the accumulation in *NPC1^I1061T/I1061T^* i^3^Neurons is less than what is observed in *NPC1^-/-^* i^3^Neurons. Consistent with an increased endolysosomal accumulation of unesterified cholesterol, EM results demonstrate that *NPC1^I1061T/I1061T^* i^3^Neurons have dark, electron-dense multilamellar structures, a well-documented feature of NPC1 disease. HPβCD has therapeutic benefits in NPC1 animal and cellular model systems [21, 22, 24, 25, 28, 29] and suggestive efficacy in NPC1 individuals in clinical trials [26]. In our study, we found HPβCD reduced cholesterol accumulation in *NPC1^I1061T/I1061T^* cells similar to the reduction demonstrated in *NPC1^-/-^* i^3^Neurons. In addition, treatment with the chaperone mo56HC improves the levels of NPC1 protein obtained from *NPC1^I1061T/I1061T^* i^3^Neurons. Thus, we confirmed that *NPC1^I1061T/I1061T^* i^3^Neurons can serve as a prototype for identifying potential therapeutic drug targets in a high throughput screen.

This study highlights the utility of an iPSC-derived neuronal model for NPC1 disease which generates a large number of homogenous neurons, provides isogenic control and replicates characteristic features of NPC1 disease caused by missense variants. This cell line provides a relevant model for researchers working on NPC1 disease and complements the existing *NPC1^-/-^*i^3^Neuron line. The added benefit of the *NPC1^I1061T/I1061T^* line comes in its utility for differential screening to identify drugs that either increase expression or stabilize the mutant NPC1 protein.

## Acknowledgments

We would like to thank Dr. Brian Blagg (University of Notre Dame, Notre Dame, IN) for the mo56HC. Help with the Sanger sequencing (Dr. Fabio Faucz, Molecular Genomics Core, NICHD) and the electron microscopy (Mr. Louis (Chip) Dye, Microscopy and Imaging Core, NICHD) is greatly appreciated. We also would like to thank Aileen Barnes for help with editing the manuscript.

## CRediT authorship contribution statement

Shikha Salhotra: Writing- review and editing, Writing-original draft, Validation, Methodology, Investigation, Formal analysis, Data Curation. Niamh X. Cawley: Writing- review and editing, Writing- original draft, Formal analysis, Data curation. Christian White: Methodology, Investigation, Analysis. Insung Kang: Methodology, Investigation, Formal analysis. Anika Prabhu: Methodology, Investigation. Cristin D. Davidson: Writing- review and editing, Formal analysis. Forbes D Porter: Writing- review and editing, Supervision, Project administration, Funding acquisition, Conceptualization, Formal analysis.

## Data availability

Data not included in the manuscript or supplemental material is available upon request.

