## Supplemental Figures for "Generation and characterization of human iPSC-derived *NPC1^I1061T/I10161T^* i^3^Neurons as a model for NPC1 disease"

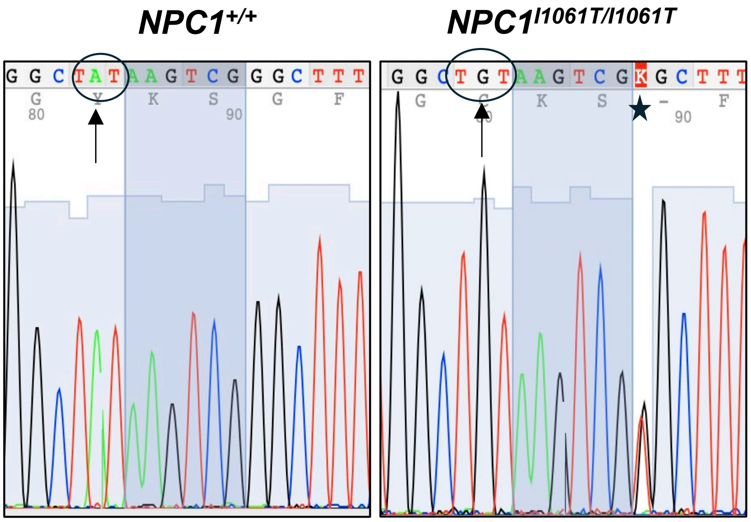


**Supplementary Figure 1.** Reverse Sanger sequencing analysis of the genomic DNA from *NPC1*$^{+/+}$ and *NPC1*$^{I1061T/I1061T}$ cells. In this direction, the A>G mutation (arrows), resulting in the isoleucine to threonine change is confirmed, along with the silent mutation (star).

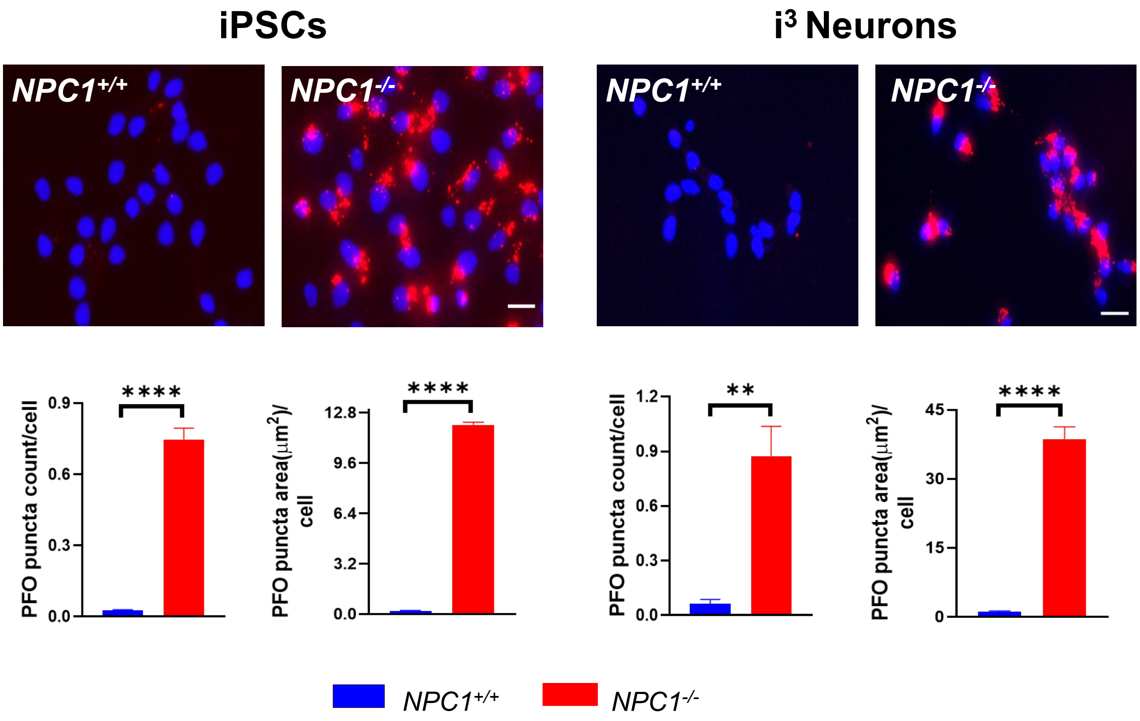


**Supplementary Figure 2.** Analysis of cholesterol accumulation in lysosomes of *NPC1*$^{+/+}$ and *NPC1*$^{-/-}$ iPSCs and i$^3$Neurons using PFO-647 staining (red). The upper panels show strong PFO staining in both *NPC1*$^{-/-}$ iPSCs and i$^3$Neurons. The lower panel shows the results of the quantification by ImageJ. The size of puncta (μm$^2$) per cell increased by 54-fold ($p<0.0001$) and 34-fold ($p<0.0001$) in *NPC1*$^{-/-}$ iPSCs and i$^3$Neurons, respectively, compared to *NPC1*$^{+/+}$ cells. Similarly, the number of PFO puncta per cell was 28-fold higher in *NPC1*$^{-/-}$ iPSCs ($p<0.0001$) and 14-fold higher in *NPC1*$^{-/-}$ i$^3$Neurons ($p<0.01$) when compared to *NPC1*$^{+/+}$ iPSCs and i$^3$Neurons, respectively. The nuclei were counterstained with DAPI (blue). 100 cells were counted per cell line and data obtained from three independent experiments (n=3). Scale bar, 20 μm.**p<0.01 and ****p<0.0001 using unpaired t-test when comparing two independent samples.

Salhotra et al. Supplementary Figure 2

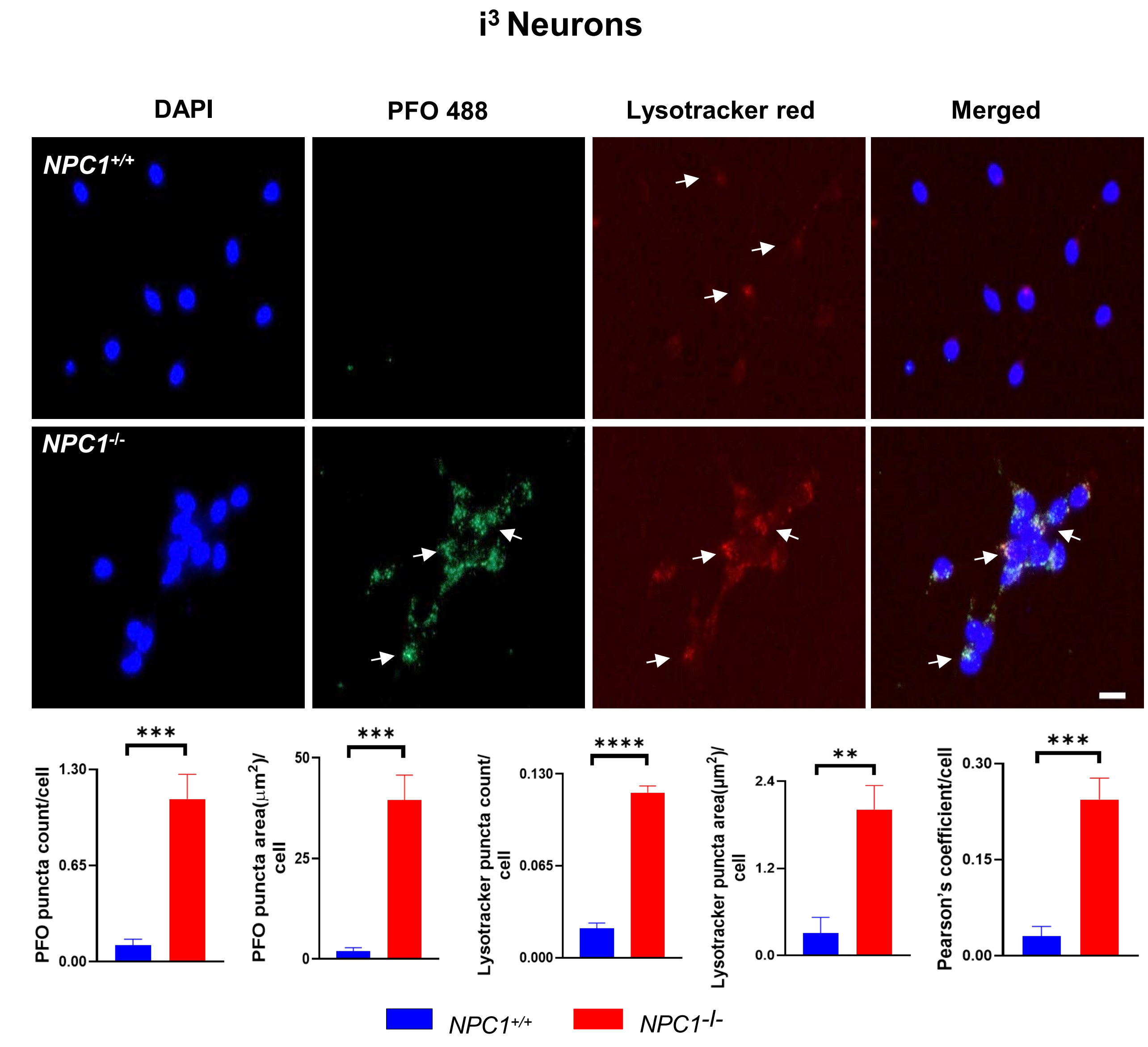


**Supplementary Figure 3.** Analysis of *NPC1*$^{+/+}$ and *NPC1*$^{-/-}$ i³Neurons for cholesterol staining with PFO-488. Unesterified cholesterol (arrows) was found to strongly co-stain with lysotracker red, indicating the accumulation in lysosomes in *NPC1*$^{-/-}$ neurons. PFO puncta area increased by 20 fold ($p<0.001$) and PFO puncta count increased by 9.8 fold ($p<0.001$) in *NPC1*$^{-/-}$ i³Neurons compared to *NPC1*$^{+/+}$. Similarly, lysotracker red puncta count and puncta area per cell increased by 5.63 fold ($p<0.0001$) and 6.5 fold ($p<0.01$), respectively in *NPC1*$^{-/-}$ i³Neurons. Pearson's coefficient was found to be 8.1 fold higher in *NPC1*$^{-/-}$ i³Neurons ($p<0.001$). The nuclei were counterstained with DAPI (blue). 100 cells were counted per cell line and data obtained from three independent experiments (n=3). Scale bar, 20 μm.**$p<0.01$, ***$p<0.001$ and ****$p<0.0001$ using unpaired t-test when comparing two independent samples.

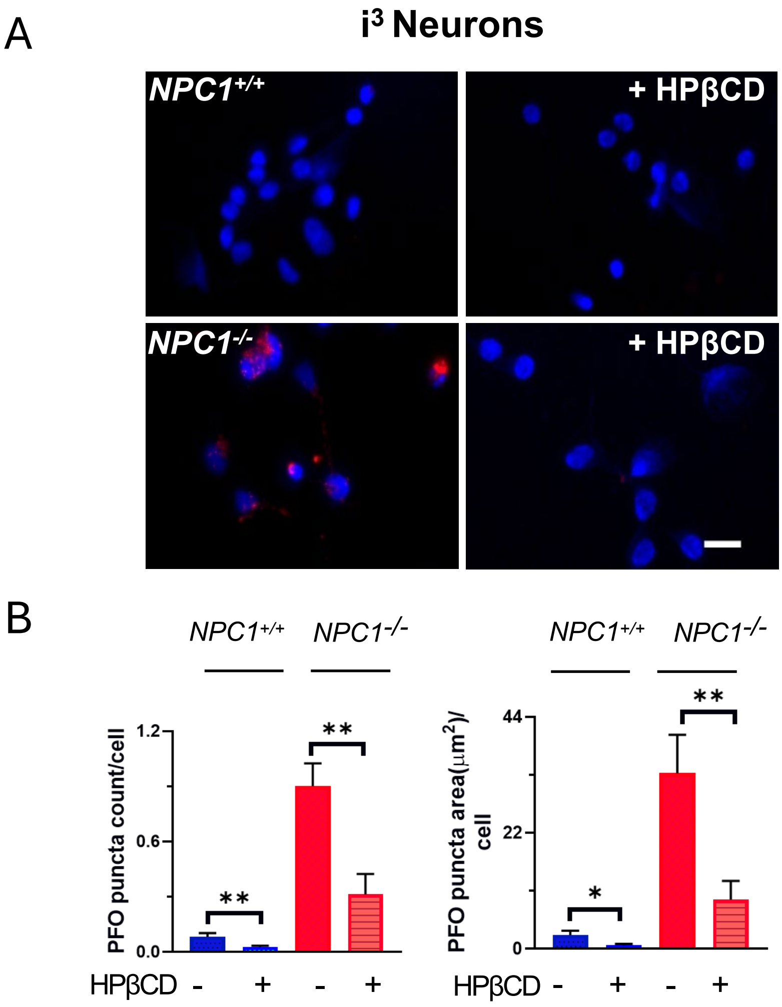


**Supplementary Figure 4.** Demonstration of the reversal of PFO staining in *NPC1$^{-/-}$* i$^3$Neurons with 2-Hydroxypropyl-β-cyclodextrin (HPβCD) treatment. (**A**) Fluorescence microscopy of *NPC1$^{+/+}$* and *NPC1$^{-/-}$* i$^3$Neurons showing PFO staining (red) in the mutant cells that is reduced when treated with HPβCD. (**B**) Quantification of the staining using ImageJ. PFO puncta count was reduced 2.9-fold ($p < 0.01$) and puncta area was reduced 3.5-fold ($p < 0.01$) after HPβCD treatment in *NPC1$^{-/-}$* i$^3$Neurons. DAPI (blue) was used to counterstain the nuclei. 100 cells were counted per cell line and data obtained from three independent experiments (n=3). Scale bar, 20 µm.*$p<0.05$, **$p<0.01$ using unpaired t-test when comparing two independent samples.
